# Role of RANKL in Alveolar Epithelial Cell Regeneration: Potential Novel Contributor to Lung Tissue Repair

**DOI:** 10.1101/2023.10.11.561640

**Authors:** Habibie Habibie, Shanshan Song, Carian E Boorsma, Kurnia S.S. Putri, Catharina Reker-Smit, Jelmer Vlasma, Mitchel J.R. Ruigrok, Robbert H Cool, Xinhui Wu, Yizhou Wang, Wim Quax, Peter Olinga, Corry-Anke Brandsma, Wim Timens, Janette Burgess, Barbro N Melgert

## Abstract

Receptor activator for NF-κβ (RANK) ligand (RANKL) is found in lung tissue and elevated in lung diseases like chronic obstructive pulmonary disease (COPD), cystic fibrosis and silica-induced pulmonary fibrosis. RANKL is a well-known stimulator of bone tissue degradation, which may explain the association between these lung diseases and osteoporosis. However, RANKL is also reported to be involved in epithelial cell regeneration in breast and thymus. We hypothesized that RANKL, which is produced directly in lung tissue, is involved in the regeneration of lung epithelial cells. Therefore, we aimed to clarify the specific role of RANKL in this process.

Using an organoid model of lung epithelial development by co-culturing primary EpCAM+ lung epithelial cells with fibroblasts, we found higher numbers of alveolar organoids after soluble RANKL treatment compared to control. Importantly, this effect was similar in human RANKL-treated organoids derived from epithelial cells isolated from lung tissue of COPD patients. The effect of RANKL was abrogated upon addition of osteoprotegerin, the soluble inhibitor of RANKL. We also found that RANKL stimulated phosphorylation of Akt suggesting involvement of its receptor RANK in the signaling pathway. Moreover, *in vivo* RANKL administration resulted in more type II alveolar epithelial cells in lungs of mice with silica-induced pulmonary fibrosis.

In conclusion, we found that RANKL promotes type II alveolar epithelial cell regeneration and may therefore be a novel contributor to lung tissue repair.

**New and Noteworthy:** Our study provides compelling evidence demonstrating an as of yet unknown function of receptor activator for NF-κβ ligand (RANKL) in lung tissue regeneration. We found that RANKL plays a role in the regeneration of lung epithelial cells, particularly type II alveolar epithelial cells. This may also have clinical implications as promotion of alveolar epithelial cell regeneration may enhance lung tissue repair, an important target in patients with lung diseases like COPD and fibrosis.

## INTRODUCTION

Receptor activator for NF-κβ (RANK) ligand (RANKL), a member of the tumor necrosis factor (TNF) superfamily, is prominently expressed in bone tissue. It is best known for its action as a stimulator of bone tissue degradation and has an important role in bone extracellular matrix regulation (1, 2). RANKL is also found in lung tissue (3) and higher levels have been reported in lung diseases like chronic obstructive pulmonary disease (COPD) (4), cystic fibrosis (5) and silica-induced pulmonary fibrosis (6). These higher levels of RANKL may explain the association of these diseases with osteoporosis (4, 5, 7). However, the lung-specific production of RANKL suggests a role in lung tissue but this role is unknown. Interestingly, RANKL has also been reported to stimulate epithelial cell regeneration in breast and thymus (8–10), which may point at a role in epithelial repair in the lung too.

Damage to lung epithelial cells is believed to be a major event for the initiation of lung diseases and it is crucial that the area of injury is quickly repaired and replaced by new epithelial cells in order to preserve the functionality of the lung (11). In alveoli, alveolar type II (ATII) epithelial cells have been shown to be a key cell type in lung repair (12). These cells can proliferate and differentiate into type I epithelial cells to regenerate the site of damage. Tissue injury will trigger secretion of factors by tissue-resident cells, that contribute to repair mechanisms including promotion of lung epithelial cells regeneration (13, 14)

In our recent study we reported that RANKL is produced directly in lung tissue by lung fibroblasts (15). We therefore postulated that RANKL expression by fibroblasts may reflect lung repair via promotion of epithelial cell regeneration to counteract the damage to epithelial cells that characterizes many lung diseases. We investigated this hypothesis using mouse and human organoids, a murine model of silica-induced pulmonary fibrosis, and epithelial cell lines.

## MATERIAL AND METHODS

### Production and purification of RANKL protein

Recombinant sRANKL was produced using standard procedures at our facility (see supplemental data for more detailed information: https://figshare.com/s/f846cfe2211b56e8bbbf). Protein purity was checked by SDS-PAGE and RANKL concentrations were measured with a Coomassie Bradford protein assay kit (Pierce, USA) using BSA as reference and with a RANKL ELISA kit (R&D, Minneapolis, USA). Values from both assays were always in the same range and the concentration obtained from the ELISA analysis was used for further experiments. Purified RANKL is routinely tested for endotoxin contamination comparing heat inactivated RANKL with native RANKL using RAW 264.7 macrophages and no measurable contamination was found.

### Animal experiments

Mice were obtained from Harlan (Horst, The Netherlands) and were maintained with permanent access to food and water in a temperature-controlled environment with a 12h dark/light cycle regimen. All experiments were approved by the Institutional Animal Care and Use Committee. Male C57BL/6 mice were used in experiments with silica-induced pulmonary fibrosis (License number DEC6064). Female and male C57BL/6N mice were used for the preparation of murine organoids (License number AVD10500202011285). Mouse experiments were performed in the animal facility of the University of Groningen according to strict governmental and international guidelines on animal experimentation.

### Immunohistochemistry

Immunohistochemical analysis of pro-surfactant protein C (SFTPC) expression was performed on 3 µm-sections of paraffin embedded zinc-fixed mouse lung tissue. Sections were deparaffinized with xylene and rehydrated with PBS. Antigen-retrieval for SFTPC was performed by incubating sections in 0.1M Tris-HCL (pH=9.0). Hereafter, an antibody against SFTPC (1:10000, MERCK) was applied for 1 h at room temperature. Afterwards, the sections were washed with PBS three times and further incubated with secondary antibody, a goat anti-rabbit IgG HRP-conjugated (DAKO) for 30 min at room temperature. Sections were further incubated with third antibody, a rabbit anti-goat IgG HRP-conjugated (DAKO) for 30 min at room temperature. Staining was visualized by NovaRED (Vector Laboratories, Burlingame, California, US). Subsequently, the tissue was counterstained with hematoxylin (Klinipath BV). After dehydration, sections were covered withDepex (SERVA, Germany) and a coverslip. Slides were scanned using a C9600 NanoZoomer (Hamamatsu Photonics, Japan) and SFTPC positive cells were quantified using ImageScope software (Leica Biosystems, Aperio, USA) by quantifying area with strong positive staining over total area of lung tissue.

### Cell lines and culture conditions

A549 epithelial cells were cultured in DMEM (GibcoTM), supplemented with 10% fetal bovine serum, 100 U/ml penicillin/streptomycin (GibcoTM) at 37°C with 5% CO2 in humidified air. CCL-206 murine lung fibroblasts were maintained in DMEM (GibcoTM) / Ham’s F12 medium (Lonza) supplemented with 10% fetal bovine serum (FBS), penicillin/streptomycin (100 U/ml, GibcoTM) and glutamine (1%, Life Technologies). MRC5 human lung fibroblasts were maintained in Ham’s F12 medium supplemented with 10% FBS, penicillin/streptomycin (100 U/ml) and glutamine (1%). Prior to organoid culture, CCL206 or MRC5 fibroblast growth was inactivated with mitomycin C (10 μg/ml, Sigma-Aldrich) for 2 h, followed by washing with PBS (GibcoTM) three times and trypsinization.

### Western Blot

A549 cells were treated with 10 ng/ml RANKL for 15 mins, 30 mins, or 2h. Cell lysates were made as previously described (16) and loaded onto a 10% Bis-Tris gel. Proteins were separated at 100V and subsequently transferred to polyvinylidene fluoride (PVDF) membranes. The membranes were blocked with 5% nonfat dry milk and incubated overnight at 4 °C with one of the following primary antibodies: Akt (1:1000 Cell Signalling), phospho-Akt (1:1000 Cell Signalling) and β-actin (1:10000 Santacruz). The membranes were further incubated with a goat anti-rabbit HRP-conjugated secondary antibody (1:2000, DAKO) for 1 h at room temperature. For detection, blotted proteins were visualized with an ECLTM Prime Western Blotting System (GE Healthcare). All expression levels were normalized to β-actin expression.

### Wnt/β-catenin Activity Assay

The TOP/FOP flash assay was performed based on a previously published protocol (17, 18). A549 cells (11000 cells/well) were seeded in 96-well plate. When confluent, cells were transfected with 100 ng/well of either TOP luciferase reporter plasmid or the negative control FOP plasmid using Lipofectamine™ LTX Reagent with PLUS™ Reagent (Invitrogen, Carlsbad, US) in serum-free Opti-MEM® medium (Life Technologies, Carlsbad, US). After 5 h of transfection, cells were stimulated with either vehicle, positive control CHIR99021 (2 µM), or RANKL (10ng/mL) in Opti-MEM® medium supplemented with 0.1% FBS for 24 hours. After stimulation, cells were lysed using the Bright-Glo™ Luciferase Assay System (Promega). Subsequently, luciferase activity was measured using a Synergy HTX Multi-Mode Microplate Reader (BioTek, Winooski, US). Data were collected with Gen5 software (BioTek).

### Human lung tissue

Human lung tissue was used for isolation of epithelial cells for organoid cultures. COPD patients included smoking or ex-smoking individuals with GOLD stages I-IV disease. Characteristics of patients can be found in supplemental table 1 (https://figshare.com/s/f846cfe2211b56e8bbbf). Subjects with other lung diseases such as asthma, cystic fibrosis, or interstitial lung diseases were excluded. The study protocol was consistent with the Research Code of the University Medical Center Groningen (research code UMCG (umcgresearch.org) and national ethical and professional guidelines (“Code of conduct for Health Research: Gedragscode-Gezondheidsonderzoek-2022.pdf (coreon.org). Lung tissues used in this study were derived from leftover lung material after lung surgery and are exempt from consent in compliance with applicable laws and regulations (Dutch laws: Medical Treatment Agreement Act (WGBO) art 458 / GDPR art 9/ UAVG art 24). Sections of lung tissue of each patient were stained with a standard hematoxylin and eosin staining and checked for abnormalities by a lung pathologist.

### Cell isolation and organoid culture

Primary lung epithelial cells from mice were isolated and cultured as previously described with modifications (19, 20). The lungs were filled with a dispase (BD Biosciences)/agarose (Sigma-Aldrich) mixture and digested at room temperature for 45 min and subsequently homogenized to a single cell suspension. Cells in suspension were negatively selected using a mix of mouse CD45-selecting (Miltenyi) and mouse CD31-selecting (Miltenyi) microbeads. Then, CD45-CD31-negative cells were further positively selected with mouse EpCAM-selecting microbeads (Miltenyi). EpCAM+ cells (10,000) and CCL206 fibroblasts (10,000) were resuspended in 100 μl DMEM/F12 medium containing 10% FBS diluted 1:1 with growth-factor-reduced Matrigel (Corning), and were seeded in a 24-well 0.4 μm transwell insert (Falcon).

Human primary lung epithelial cells were isolated from lung tissue of patients with COPD. Distal human lung tissue was dissociated and homogenized with a multi-tissue dissociation kit 1 (Miltenyi) using a GentleMACS Octo dissociator at 37 °C (Miltenyi). Cells in the resulting suspension were negatively selected using a mix of human CD45-selecting (Miltenyi) and human CD31-selecting (Miltenyi) microbeads. The CD45-CD31-negative cells were further positively selected with human EpCAM-selecting microbeads (Miltenyi). EpCAM+ cells (5000) were seeded with MRC5 fibroblasts (5,000) in Matrigel.

After solidifying, organoid cultures were maintained in DMEM/F12 medium with 5% (v/v) FBS, 1% insulin-transferrin-selenium (GibcoTM), recombinant mouse EGF (0·025 µg/ml, Sigma-Aldrich), and bovine pituitary extract (30 µg/ml, Sigma-Aldrich). To prevent dissociation-induced apoptosis, ROCK inhibitor (10µM, Y-27632, TOCRIS) was added for the first 48 h. Organoid cultures from mouse and human lung tissue were treated with RANKL (10ng/ml), osteoprotegerin (OPG, 40ng/mL, R&D, Minneapolis, USA), a combination of RANKL and OPG (RANKL 10ng/mL + OPG 40ng/mL incubated 37 °C for 1 hour before addition) or vehicle. All treatments were freshly added to the cultures every two days. Organoid cultures were maintained at 37 °C with 5% CO2 in humidified air. The number of organoids was manually counted and organoid diameter was measured at day 14 of the culture using an Olympus IX50 light microscope connected to Cell^A software (Olympus).

### Immunofluorescence analysis

Lung organoids were stained for acetylated α-tubulin and pro-surfactant protein C to identify ciliated airway epithelial cells and ATII cells respectively, as described previously (19, 20). In short, organoids were fixed with ice-cold acetone/methanol (1:1 v/v) and a specific antibody binding was blocked with 5% BSA (w/v, Sigma-Aldrich). Organoids were then incubated with primary antibodies against acetylated α-tubulin (Santa Cruz Biotechnology) and SFTPC (MERCK) in a 1:200 dilution in PBS with 0·1% (w/v) BSA and 0·1% (v/v) Triton-X100 (Thermo Fisher) at 4 ℃ overnight. Organoids were washed three times with PBS and incubated with the secondary antibodies (1:200, donkey anti-rabbit Alexafluor 488, Invitrogen; 1:200, donkey anti-mouse Alexafluor 568, Invitrogen) for 2 h at room temperature. Organoids were washed with PBS three times again and then cut from inserts and transferred onto glass slides with mounting medium containing DAPI (Abcam) and a coverslip. Images were obtained using a Leica DM4000b fluorescence microscope connected to Leica Application Suite software.

### Silica-induced lung fibrosis in mice

Fibrosis was induced using Min-U-Sil 5 crystalline silica (a kind gift from Dr. Andy Ghio, US EPA, Chapel Hill, NC) by a method described previously (21, 22). On day zero animals were anesthetized using isoflurane before they received a single administration of crystalline silica (0.2 g/kg) in 50 μl 0.9% saline by intratracheal installation, while control animals received an equivalent volume of 0.9% saline. After 28 days, RANKL treatment began and was continued for an additional 2 weeks. We calculated that the administered RANKL dose should exceed the calculated OPG content (0.8 mg OPG/lung) (22, 23) in fibrotic lungs to be able to see a biological effect of RANKL. We therefore used doses of 5 μg and 10 μg RANKL per mouse three times per week. RANKL was administered intranasally in 40 μl saline. Control mice with pulmonary fibrosis received 40 μl saline (vehicle) three times per week. On day 43, mice were sacrificed and lungs were collected for flow cytometry and histology.

### Flow cytometry

The left lungs of silica/vehicle-treated mice were used for isolation of a single cell suspension for flow cytometry (see supplemental data for more detailed information: https://figshare.com/s/f846cfe2211b56e8bbbf). Total cell numbers were determined using a FACS Array cell counter (BD Biosciences) and used to calculate total numbers of specific cell types. Lung single cell suspensions were stained for epithelial cells and RANKL expression by flow cytometry (see supplemental Table 2 for antibody details, https://figshare.com/s/f846cfe2211b56e8bbbf). Samples were analyzed using an LSRII flow cytometer (BD Biosciences). Data was analyzed using FlowJo software (Tree Start, Ashland, USA). See supplemental figure 1 for the gating strategy (https://figshare.com/s/f846cfe2211b56e8bbbf).

### Statistical analyses

Data sets with n ≤8 were not considered to be normally distributed. Statistical differences between two groups were assessed using nonparametric Mann-Whitney U for unpaired data or Wilcoxon for paired data. For the comparison of multiple groups, we used Friedman or Kruskal Wallis tests. p<0.05 was considered significant. The data were analyzed using GraphPad Prism 8 (GraphPad software, San Diego, USA).

## RESULTS

### RANKL promotes growth of murine and human lung organoids and this effect is inhibited by osteoprotegerin (OPG)

To assess if RANKL treatment has effects on primary epithelial cells, we investigated RANKL treatment of lung organoids generated from both human and murine epithelial cells. Murine organoids will grow into structures resembling airways and alveoli and the distinct morphologies are shown in fig. 1a. We confirmed these two different phenotypes by immunofluorescence staining using acetylated alpha-tubulin (marker of ciliated airway epithelium) and pro-surfactant protein C (marker of ATII epithelium) (fig. 1b).

**Figure 1.**
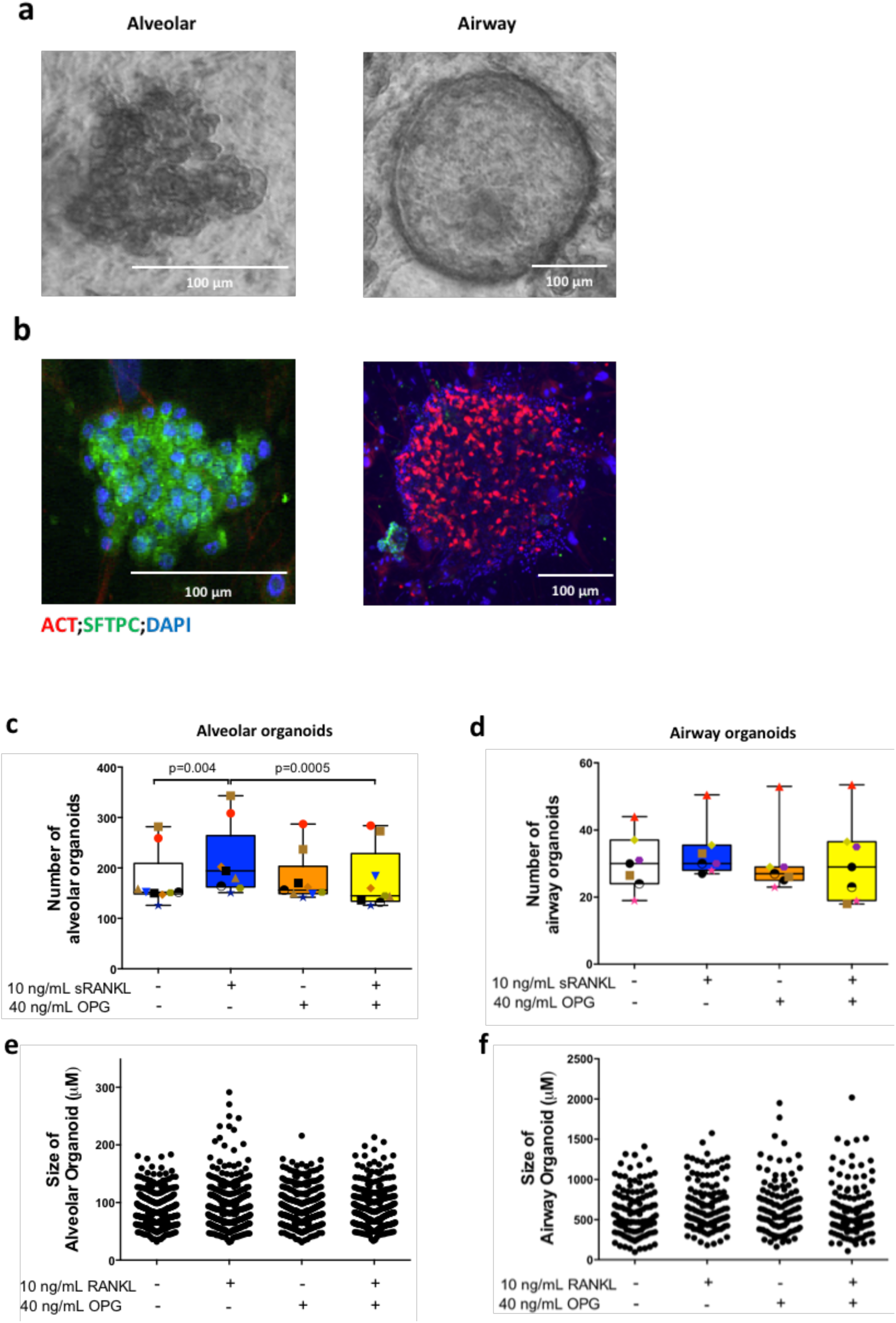
RANKL promotes growth of murine alveolar organoids and this effect is inhibited by osteoprotegerin (OPG). (a) Light microscopy images of murine alveolar and airway organoids. (b) Immunofluorescence images of murine alveolar and airway organoids stained for acetylated tubulin (ACT, red), pro-surfactant protein C (SFTPC, green), and nuclei (DAPI, blue). Quantification of number (c-d, each donor marked with different color and symbol) and size (e-f) of murine alveolar organoids and airway organoids after 14 days of incubation with either RANKL (10ng/mL), OPG (40 ng/mL), the combination of RANKL and OPG, or vehicle. Groups were compared using a Friedman multiple comparison test, p<0.05 was considered significant.

RANKL treatment resulted in significantly more alveolar organoids as compared to untreated controls and no effect was seen on airway organoids (fig. 1c and 1d). Interestingly, co-treatment with osteoprotegerin, the soluble inhibitor of RANKL, abrogated the stimulatory effect of RANKL on alveolar organoids (Fig. 1c and 1d). RANKL treatment did not have an effect on the size of either alveolar or airway organoids (Fig.1e and 1f).

To investigate whether these findings in mice could be replicated with human epithelial cells, we also generated organoids from lung epithelial cells obtained from patients with COPD having lung reduction surgery as a treatment for severe emphysema or suspected tumors. These human organoids do not develop into two distinct morphologies as seen for murine organoids, but have a mixed phenotype containing both alveolar and airway epithelial cells (Fig. 2a-b). Similar to murine organoids, RANKL treatment also significantly promoted formation of human lung organoids, and this effect was also inhibited by OPG (Fig. 2c). The size of organoids was again not affected by the presence of RANKL (Fig. 2d). Taken together, these observations imply that RANKL promotes growth of both primary murine and human ATII cells.

**Figure 2.**
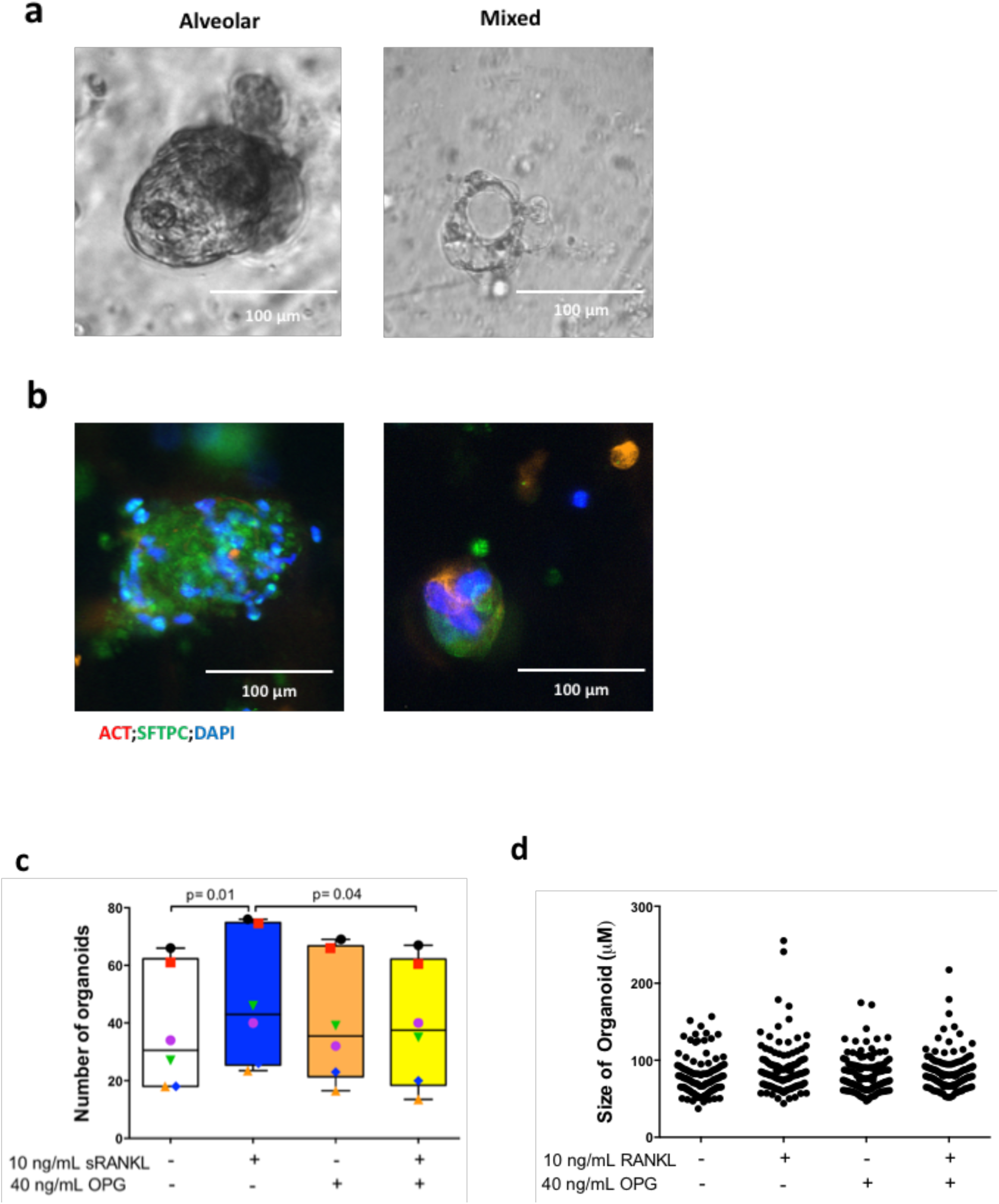
RANKL promotes growth of human lung organoids and this effect is inhibited by osteoprotegerin (OPG). (a) Light microscopy image of human lung organoids. (b) immunofluorescence image of human lung organoids stained for acetylated tubulin (ACT, red), pro-surfactant protein C (SFTPC, green), and nuclei (DAPI, blue). Quantification of the number (c, each donor marked with different color and symbol) and size (d) of human lung organoids after 14 days of incubation with either RANKL (10ng/mL), OPG (40 ng/mL), the combination of RANKL and OPG, or vehicle. Groups were compared using a Friedman multiple comparison test, p<0.05 was considered significant.

**Figure 3.**
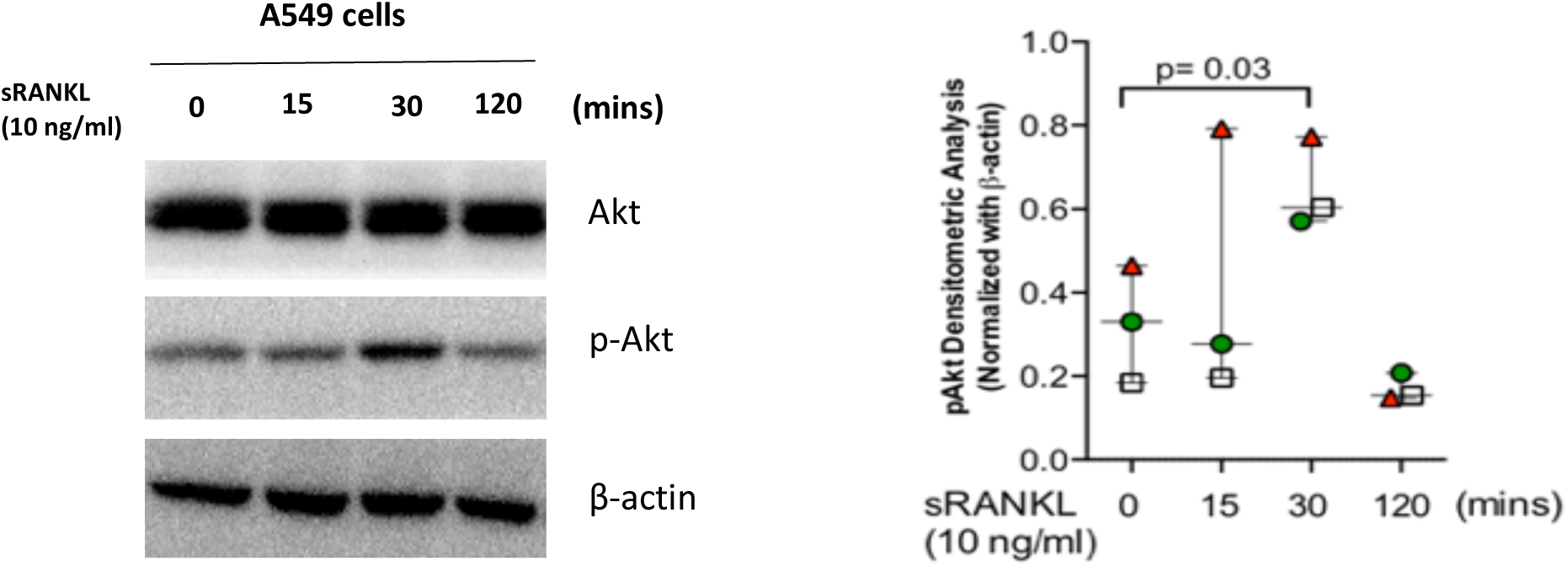
RANKL triggers activation of the Akt signaling pathway in A549 epithelial cells. A549 epithelial cells were cultured with 10 ng/mL RANKL and cell lysates were analyzed for phosphorylation of Akt (p-Akt) at several time points by Western blot (n=3). Beta-actin was used as a loading control. Groups were compared using a repeated measures ANOVA, p<0.05 was considered significant.

### RANKL triggers activation of Akt signaling pathway in alveolar type II epithelial cells

To elucidate which receptor is involved in RANKL-induced ATII cell growth, we investigated signaling pathways related to RANK and Leucine Rich Repeat Containing G Protein-Coupled Receptor 4 (LGR4), both being reported receptors for RANKL (24). RANK was previously shown to signal through the Akt pathway, while LGR4 was reported to use the WNT/ beta-catenin pathway (25). We stimulated A549 epithelial cells with RANKL and analyzed phosphorylation of Akt at several time points by Western blotting and beta-catenin transcriptional activity with a TOP/FOP dual-luciferase reporter system. Stimulation with RANKL resulted in significant phosphorylation of Akt at 30 minutes after the start of the RANKL stimulation (fig.3 and supplemental fig.2 for all original western blot data, https://figshare.com/s/f846cfe2211b56e8bbbf), while no activation of beta-catenin was found (supplemental fig.3, https://figshare.com/s/f846cfe2211b56e8bbbf). This suggests that RANKL induces ATII proliferation through an interaction with RANK.

### Intranasal RANKL treatment of mice with silica-induced pulmonary fibrosis promotes epithelial cell number

To assess the effect of RANKL on lung epithelial cell growth in vivo in a condition known to have impaired epithelial repair (26), we treated mice with silica-induced pulmonary fibrosis with two different doses of recombinant RANKL. As expected, induction of pulmonary fibrosis resulted in fewer EpCAM+ cells in lung tissue as compared to sham-treated control mice (Fig 4a). Treatment with 5 ug intranasal RANKL treatment normalized this loss of EpCAM+ cells in fibrotic lung tissue as compared to saline-treated mice and treatment with 10 ug resulted in even higher numbers of EpCAM+ cells (fig. 4a). This higher number of epithelial cells was paralleled by a similar pattern for the number of RANKL and EpCAM double-positive cells (fig. 4b). To further investigate the identity of these EpCAM+ cells, tissue sections were stained for pro-surfactant protein C (fig. 4c). RANKL administration resulted in significantly more area staining positively for pro-surfactant protein C in lung tissue (fig. 4d). Taken together, these observations imply that RANKL promotes regeneration of ATII cells in mice.

**Figure 4.**
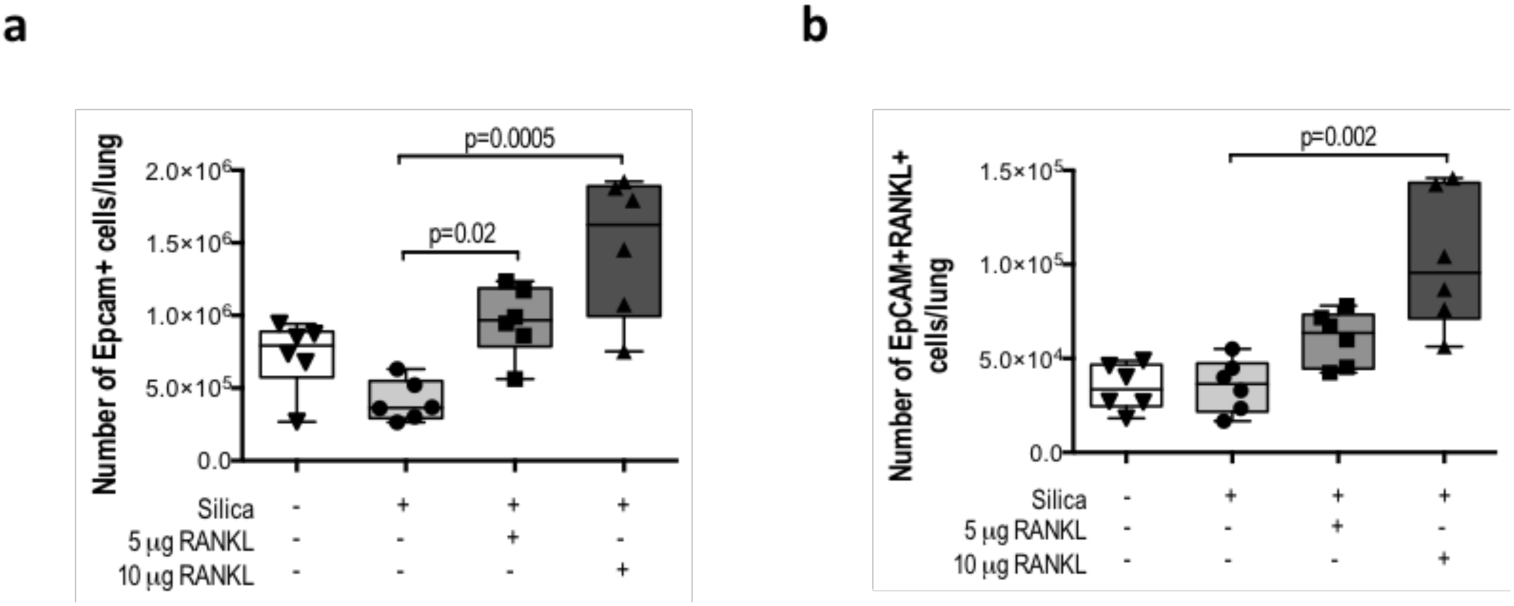
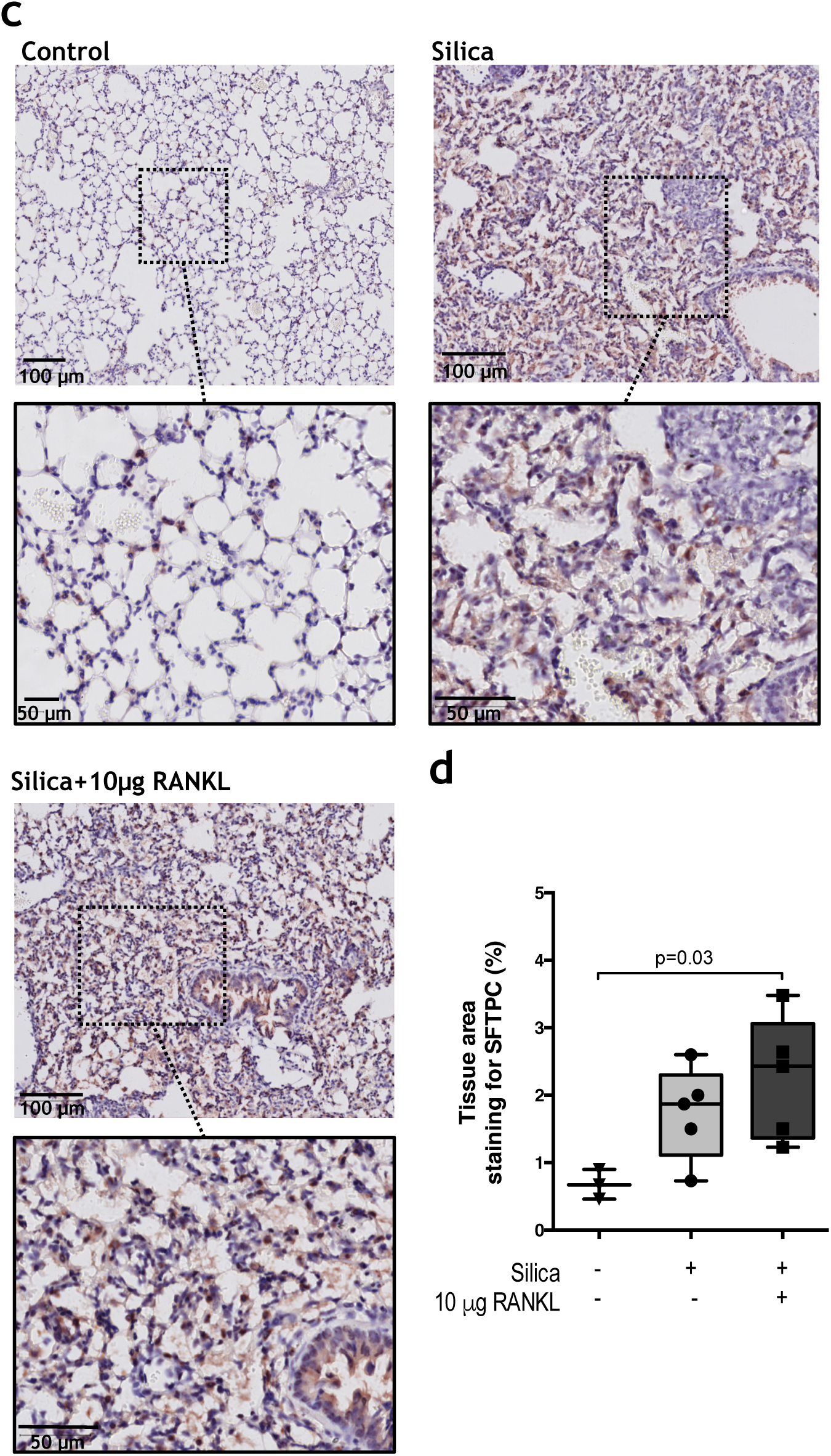
Intranasal RANKL treatment of mice with silica induced pulmonary fibrosis promotes epithelial cell proliferation. Mice were treated with RANKL (3x per week for two weeks, 5 or 10 µg per mouse/administration) four weeks after inducing pulmonary fibrosis with silica. (a) EpCAM+ epithelial cells and (b) EpCAM-RANKL double-positive epithelial cells were assessed by flow cytometry. (c) Representative sections of lung tissue from vehicle-treated, silica-treated and silica+10 µg RANKL-treated mice stained for pro-surfactant protein C (SFPTC: red, nuclei: blue). (d) Quantification for pro-surfactant protein C staining. Differences between groups were tested with a Kruskall Wallis test, p<0.05 was considered significant.

## DISCUSSION

Our study has shown that RANKL contributes to lung epithelial cell regeneration. Moreover, our data reveal that RANKL contributes specifically to an increase of ATII-containing organoids from both mouse and human lung tissue. ATII cells function as progenitors for new ATII and ATI cells (27, 28) and have been shown to be a crucial factor in the lung repair and regeneration (12, 13). This exciting novel function of RANKL may therefore be further investigated as a therapeutic approach aiming at stimulating lung tissue repair in lung diseases characterized by insufficient epithelial repair like COPD, cystic fibrosis and pulmonary fibrosis.

RANKL has been previously identified as a key regulator of epithelial cell proliferation and differentiation in various tissues, such as the thymus (8), mammary glands (9), and epidermo-pilosebaceous units of hair follicles (29). It has also been reported that RANKL administration boosts thymic epithelial regeneration upon bone marrow transplantation (10). In combination with our data, the evidence suggests that RANKL possesses an innate ability to stimulate the growth of several epithelial cell types. Interestingly, we recently showed that RANKL confers protection against cell death in precision-cut lung slices (30). Therefore, the higher numbers of ATII cells we found in organoids and *in vivo* after RANKL treatment may be the result of improved survival and not necessarily only of induction of proliferation.

RANKL was previously reported to signal via RANK or LGR4 receptors (25) and our results suggest that RANKL in lung tissue specifically uses RANK to signal. We found that RANKL activated downstream signaling of the RANK pathway in A549 cells, which is similar to what was found for mammary gland and thymic epithelial cells (31–37). In addition, we also found that RANKL treatment stimulated phosphorylation of Akt, which is a known regulator of tissue regeneration (38) and this observation was in line with the effect of RANKL in lung slices (30). Collectively these results suggest that RANKL stimulates regeneration of lung epithelial cells through its receptor RANK.

We investigated the effects of RANKL on primary epithelial cells derived from both mouse and human lung tissue using a model of lung organoids. In murine organoids, RANKL specifically stimulated outgrowth of alveolar epithelial-type organoids but not of airway epithelial-type organoids. Although we did not observe dedicated airway epithelial-type organoids in cultures derived from human cells, RANKL treatment did also induce more outgrowth of mixed-type lung organoids suggesting that RANKL induces alveolar regeneration in both human and mouse. The reason for the lack of effect on airway epithelial cells is unclear but may be caused by different expression of RANK on these different types of epithelial cells. Data from the various datasets available on ipfcellatlas.com and copdcellatlas.com suggests that the expression of RANK mRNA (TNFRSF11a) is indeed higher in ATII cells compared to airway basal cells.

RANKL levels have been found to be elevated in patients with lung diseases such as COPD (4) and cystic fibrosis (5), as well as in BAL fluid and in ATII cells of mice with silica-induced fibrosis (6). The cellular origin of this RANKL production is not completely clear yet, but it is at least produced by lung fibroblasts as we have shown in our previous study (3, 15). These findings suggest that the production of RANKL in lung tissue and the increase during lung disease may be associated with maintaining epithelial cells via enhancing epithelial cell regeneration. We supported this notion by treating mice with silica-induced fibrosis, a known model of impaired epithelial repair (26), with RANKL and showing that this strongly enhanced the number of ATII epithelial cells in fibrotic lung tissue compared to saline treatment. Although we did not treat healthy mice with RANKL to show an effect of RANKL on healthy lung tissue, our experiment did support that in condition of epithelial damage, RANKL can contribute to epithelial proliferation, which is when it is needed most. In addition, recent work by Ju *et al* showed that a RANKL partial peptide could protect again bleomycin induced pulmonary fibrosis, suggesting there may be additional benefits to using some of RANKL’s function for pulmonary fibrosis (39). Together, these data suggest that RANKL may be used as a therapeutic strategy for stimulating alveolar repair in diseases with impaired alveolar repair, like IPF or COPD, though this needs to be tested further in dedicated models.

In conclusion, we found that RANKL promotes ATII cell regeneration and may therefore contribute to alveolar repair. Our data suggest that the higher levels of RANKL found in several lung diseases may reflect stimulation of epithelial repair by the lung to counteract the lung tissue destruction that characterizes these lung diseases.

## ACKNOWLEDGEMENT

The funding for this research was partly provided by the Indonesian Endowment Fund for Education (LPDP Scholarship), which supported H.H in pursuing his Ph.D studies at the Groningen Research Institute of Pharmacy, University of Groningen in the Netherlands.

## SUPPLEMENTAL DATA

https://figshare.com/s/f846cfe2211b56e8bbbf

## Notes

https://figshare.com/s/f846cfe2211b56e8bbbf

### Competing Interest Statement

The authors have declared no competing interest.

